# Multi-omics data analysis supporting inheritance of acquired traits in humans

**DOI:** 10.1101/2023.05.10.540125

**Authors:** Abhay Sharma

## Abstract

Experimental evidence supports presence of germline mediated nongenetic inheritance in animals including mammalian models. Observational and molecular epidemiological studies also suggest existence of environmental exposure induced germline inheritance in humans. Considering the obvious difficulties in conducting prospective cross-generational human studies, integrative analysis of available multi-omics data may seem to offer a valuable approach for assessing biological plausibility of inheritance of acquired traits in the species. To that end, the concept has mainly been tested here that, under the assumption of inheritance, human exposure to an environmental factor, if known to induce paternal inheritance in mammalian models and to act through molecular pathways conserved across species, would induce correlated changes in sperm epigenome and somatic transcriptome in the exposed subjects. The test is based on physical exercise, a physiologically relevant factor for which multiple datasets of human sperm DNA methylome and somatic transcriptome from interventional pre-post cohort studies, that minimizes inter-individual variability, were available. Existing somatic transcriptome datasets from animal model studies on exercise have been used for comparison. To control the analysis, bariatric surgery, for which exercise matched human datasets existed, have been used. The hypothesis testing involves gene set overrepresentation, comparison between directionality of epigenome and coding transcriptome changes, gene ontology enrichment, and epigenome and non-coding transcriptome interaction. Remarkably, the results show that, in humans, exercise induced DNA methylation changes in sperm specifically represent transcriptional response to exercise in soma. This germline epigenomic encoding of acquired transcriptome changes in soma clearly supports biological plausibility of epigenetic inheritance in humans.

## Introduction

Accumulating evidence supports presence of nongenetic germline inheritance of induced traits in animals including mammals (Anastasiadi *et al*., 2021; Beil *et al*., 2023; Cavalli and Heard, 2019; Dali et al., 2023; Fitz-James and Cavalli, 2022; Perez and Lehner, 2019; Santilli and Boskovic, 2023; Tando and Matsui, 2023). This unconventional, non-DNA sequence-based transmission of phenotypic information across organismal generations, often referred to as transgenerational epigenetic inheritance (TEI), is considered to be mediated by factors such as DNA methylation, histone post-translational modifications, and RNA (Cavalli and Heard, 2019; Perez and Lehner, 2019; Anastasiadi *et al*., 2021; Santilli and Boskovic, 2023; Takahashi *et al*., 2023). Besides TEI, intergenerational inheritance, that may not necessarily implicate germline transmission but nonetheless share features with the former, is also considered relevant in nongenetic inheritance. Overall, germline mediated epigenetic inheritance contradicts multiple fundamental concepts in biology including neo-Darwinian principle that limits the source of heredity exclusively to DNA sequence, Weismann barrier that prohibits soma to germline transfer of hereditary information, and developmental reprogramming that inhibits germline inheritance of epigenetic signals, and hence has intriguing implications for ecology, evolution, and health and disease (Cavalli and Heard, 2019; Anastasiadi *et al*., 2021; Fitz-James and Cavalli, 2022; Santilli and Boskovic, 2023; Adrian-Kalchhauser *et al*., 2020).

Environmentally induced maternal TEI, first described as a consequence of exposure of gestating female rats to an endocrine disrupting compound (Anway *et al*., 2005), has been shown in invertebrates and vertebrates alike, with triggers including toxicants and pollutants, dietary factors, and stress, besides others (Bohacek and Mansuy, 2017; Cavalli and Heard, 2019; Dali et al., 2023; Perez and Lehner, 2019; Fitz-James and Cavalli, 2022; Santilli and Boskovic, 2023; Tando and Matsui, 2023). However, difficulties in eliminating confounders like genetic differences especially in rat models lacking inbred stocks, maternal contribution, and *in utero* and postnatal effects in female exposure TEI models remain a concern (Bohacek and Mansuy, 2017; Cavalli and Heard, 2019). Given this, male exposure and patrilineal models of inheritance involving founders with similar genetic background are considered relatively less confounding and hence the focus of recent investigations (Bohacek and Mansuy, 2017; Cavalli and Heard, 2019; Perez and Lehner, 2019). With environmentally induced paternal TEI first described in the fruit fly *Drosophila* using a neuroactive compound as the trigger in an isogenic fly line (Sharma and Singh, 2009), male exposure and male line mediated inheritance, inter- or trans-generational, has been shown in various species including mammals (Beil et al., 2023; Bohacek and Mansuy, 2017; Perez and Lehner, 2019; Fitz-James and Cavalli, 2022; Santilli and Boskovic, 2023; Tando and Matsui, 2023).

Epidemiological and ancestral exposure studies also suggest potential presence of epigenetic inheritance in humans (Hart and Tadros, 2019; Senaldi and Smith-Raska, 2020; Vågerö *et al*., 2022; Dachew *et al*., 2023; Kaufman *et al*., 2023; Niu *et al*., 2023; Santilli and Boskovic, 2023). Additionally, considering the obvious difficulties in conducting human cross-generational studies (Senaldi and Smith-Raska, 2020; Santilli and Boskovic, 2023), molecular support for paternal germline inheritance of environmental effects in humans has recently been found in terms of father’s smoking showing association with offspring blood DNA methylation (Kitaba *et al*., 2023). Nevertheless, with integrative genome scale data analysis (Molaro *et al*., 2011; Cavalli and Heard, 2019; Sharma, 2020; Niu *et al*., 2023) and leveraging animal evidence (Perez and Lehner, 2019; Fitz-James and Cavalli, 2022; Bhalla and Sharma, 2023) considered as valuable approaches in epigenetic inheritance, it remains unexplored if applying these approaches could provide further insights into possible existence of nongenetic germline inheritance in humans. For example, subject to availability of data, it could be explored if sperm epigenome encodes somatic transcriptomic changes induced by a physiologically relevant environmental factor, provided the latter is known to act through molecular pathways conserved across species and to trigger paternal inheritance in mammalian models. The concept behind exploring this is that positive evidence of the above would raise the biological plausibility of epigenetic inheritance in humans. Encouragingly, as it fits the said criteria, physical exercise may serve as an ideal physiological intervention for testing the concept. First, studies employing a pre-post design, that minimizes inter-individual variability (Wang *et al*., 2005; Gallego-Paüls *et al*., 2021), for identifying human exercise associated sperm DNA methylation changes (Ingerslev *et al*., 2018) as well as somatic coding and non-coding transcriptomic alterations have been described, with a multitude of the latter available in PubMed and Gene Expression Omnibus (GEO) databases (Sayers *et al*., 2022). Second, exercise responses are similar across species including rat, mouse, and *Drosophila* (Muñoz *et al*., 2023; Watanabe and Riddle, 2019), for which multiple exercise associated somatic transcriptome datasets are also available in the databases (Sayers *et al*., 2022), allowing comparison between humans and animal models. Given that exercise induced paternal inheritance, as also maternal, is known in rodents (Kusuyama *et al*., 2020; Vieira de Sousa Neto *et al*., 2021), the hypothesis has been tested here, using the aforesaid multi-omics data, that exercise induced DNA methylation changes in human sperm specifically represent conserved transcriptional response to exercise in soma. The analysis is based on gene set overrepresentation test, comparison between regulatory directionality of epigenetic and transcriptional changes, gene ontology enrichment test, and epigenetic and non-coding transcriptome correlation. To control the analysis, bariatric surgery, for which exercise matched human cohort datasets were available (Donkin *et al*., 2016), has been used. Remarkably, the results suggest that exercise induced gene expression characteristics are indeed encoded in human sperm epigenome, thus supporting inheritance of induced traits in our species.

## Methods

### Data collection

For differentially methylated genes (DMGs) in human sperm following exercise or bariatric surgery, full sets of original author-identified genes annotating to differentially methylated regions with <0.05 adjusted *p* value were considered, irrespective of percentage of methylation difference. The DMGs showing both hyper- and hypo-methylation, depending on chromosomal regions, were not considered. Differentially expressed mRNAs (DEmRNAs) in human somatic tissues following exercise or bariatric surgery, and in rat, mouse, and *Drosophila* somatic tissues following exercise were retrieved from original reports, if provided in full, or identified from the corresponding GEO submissions, mainly using GEO2R (Barrett *et al*., 2013) and GREIN (Mahi *et al*., 2019), for microarray and RNA-seq data, in that order, using the criteria of <0.05 adjusted *p* value and >1.2 fold change. Duplicates, if any, were removed in excel after alphabetically sorting the gene symbols. Differentially expressed miRNAs (DEmiRNAs) were retrieved from original studies, using the criteria of <0.05 adjusted *p* value and >1.2 fold change. The gene symbols for DMGs and DEmRNAs were harmonized using, as appropriate, HumanMine (Lyne *et al*., 2022), rat and mouse options in MouseMine (Motenko *et al*., 2015), and FlyMine (Lyne *et al*., 2007) first, and then using Multi-symbol checker for HGNC (Seal *et al*., 2023), g:Convert for RGD and MGI (Reimand *et al*., 2007), and FlyBase ID Validator for *Drosophila* (Gramates *et al*., 2022). The miRNA nomenclature used in the original studies related to DEmiRNAs was considered as such.

### Gene set overrepresentation and other tests

Gene set overlap was examined in excel using hypergeometric test with sampling without replacement from a finite population. A population size of 25000 human genes was used in the test. The rat and mouse gene symbols in upper case were used for finding overlap with human gene sets. For overlap between *Drosophila* and human gene sets, human orthologs of fly genes identified using DIOPT Ortholog Finder (Hu *et al*., 2011), were used, irrespective of rank and scores. RANDOM.ORG (Haahr, 2023) was used to generate random numbers for gene symbol randomization. The chi-square goodness of fit test (Stangroom, 2018) was performed to compare observed and expected probability distribution.

### Gene ontology enrichment

Enrichment of gene ontology biological process terms was determined using Gene Ontology Resource (Gene Ontology Consortium, 2021), with all human genes as the default reference, and Fisher’s exact test and <0.05 FDR as statistical and >1.2 fold as enrichment criteria. For enrichment analysis, human, rat, and mouse genes were used directly, whereas, for *Drosophila* genes, human orthologs were considered.

### miRNA targets

Enrichment of miRNA target genes was found using ToppGene Suite (Chen *et al*., 2009), with all human genes as default background, and <0.05 Bonferroni *q* value as statistical significance. The miRNA targets were identified using TargetScan (Agarwal *et al*., 2015), irrespective of aggregate P_CT_ and cumulative weighted context score.

### Visualization

The DMGs, DEmRNAs, DEmiRNAs, and gene ontology enrichment heatmaps were produced in Heatmapper (Babicki *et al*., 2016) using methylation difference, log_2_ fold change, and log_2_ fold enrichment, as appropriate, with the scale type none, and, wherever applicable, the clustering method average linkage and the distance measurement method Euclidean. Bar graphs displaying log_2_ fold enrichment in gene set overlap and gene ontology enrichment, and -log_10_ *q* value in miRNA target enrichment were produced in excel.

## Results and Discussion

### Data compendium

A compendium of DMG, DEmRNA, and DEmiRNA datasets belonging to 57 genome level studies was developed for performing the present analysis. It is presented in **Supplementary Table 1**, along with percentage of methylation difference for DMGs and log_2_ fold change for DEmRNAs and DEmiRNAs, original publication and data source references, and data collection process, as also unique identifiers assigned to each of the 118 gene or miRNA set present in the compendium for describing the results here. The data in the compendium comprise of exercise associated DMGs in human sperm (Ingerslev *et al*., 2018), DEmRNAs in human skeletal muscle, the main target tissue of exercise (Vina *et al*., 2012), and whole blood and blood cells, and DEmiRNAs in human skeletal muscle, and blood cells and plasma. The control, bariatric surgery associated human datasets in the compendium include DMGs in sperm (Donkin *et al*., 2016), and DEmRNAs in subcutaneous adipose tissue. Also, exercise associated animal datasets – DEmRNAs in rat and mouse skeletal muscle, and in *Drosophila* larval body wall muscle and fly as a whole – are included.

### Analysis design

A five-step analysis was designed mainly on the basis of data available in the compendium. First, gene set overrepresentation of human exercise sperm DMGs in human exercise somatic DEmRNAs was to be examined, in order to understand if exercise induced changes in germline and soma are specifically shared in terms of affected genes. Next, gene set overrepresentation of human exercise sperm DMGs in animal exercise somatic DEmRNAs was to be tested, to find if the above sharing of genes relates to exercise response conserved across species. This was to be followed by gene ontology analysis for enrichment of biological process terms in the gene sets representing human exercise sperm DMGs, and human and animal somatic DEmRNAs, for confirming that the DMGs and the DEmRNAs relate to processes relevant in exercise biology. Hereafter, the analysis was to focus on human data alone. As promoter methylation is generally considered to inversely regulate gene expression (Pepin *et al*., 2019), regulatory directionality of exercise associated sperm promoter DMGs and somatic DEmRNAs were to be compared next, for examining if an inverse relationship exists between the two, and thereby germline epigenetic encoding of somatic traits is supported. Finally, given that miRNAs are considered as essential mediators of exercise associated processes (Silva *et al*., 2017), the analysis was to focus on miRNA target enrichment in exercise sperm DMGs, exercise associated somatic DEmiRNAs, and gene ontology enrichment in the targets of these DEmiRNAs, for additionally examining representation of exercise biology in sperm epigenome. Positive evidence in each of the above steps of analysis were to support the concept that exercise induced changes in human sperm DNA methylation represent conserved transcriptional response to exercise in soma in specific, and exercise biology in general.

### Overlap between human exercise sperm DMGs and human exercise somatic DEmRNAs

A gene set overlap test was first performed to examine if exercise sperm DMGs specifically overrepresent exercise somatic DEmRNAs in humans. Independent clustering of exercise associated sperm DMGs, and skeletal muscle and blood DEmRNAs, and bariatric surgery associated sperm DMGs and subcutaneous adipose DEmRNAs showed largely similar regulation between samples within each category (**Fig. 1a-e**), and hence the genes in each of these categories were separately pooled for the test, to achieve higher statistical strength with increased gene set size. To avoid size related bias in overlap tests (Wang *et al.,* 2010), bariatric surgery sperm DMGs, that was almost 10 times higher in number in comparison to exercise DMGs, were randomized and divided into 10 size matched subsets. A total of 11 sperm DMG lists, one related to exercise and the rest to bariatric surgery, were thus finally tested for overlap with the pooled DEmRNA gene sets. Remarkably, the exercise sperm DMGs, compared to bariatric surgery sperm DMGs, showed unequivocally higher enrichment of exercise, not bariatric surgery, related DEmRNAs (**Fig. 1f**). These results demonstrated that exercise sperm DMGs and exercise somatic DEmRNAs are specifically related in terms of affected genes.

**Fig. 1.**
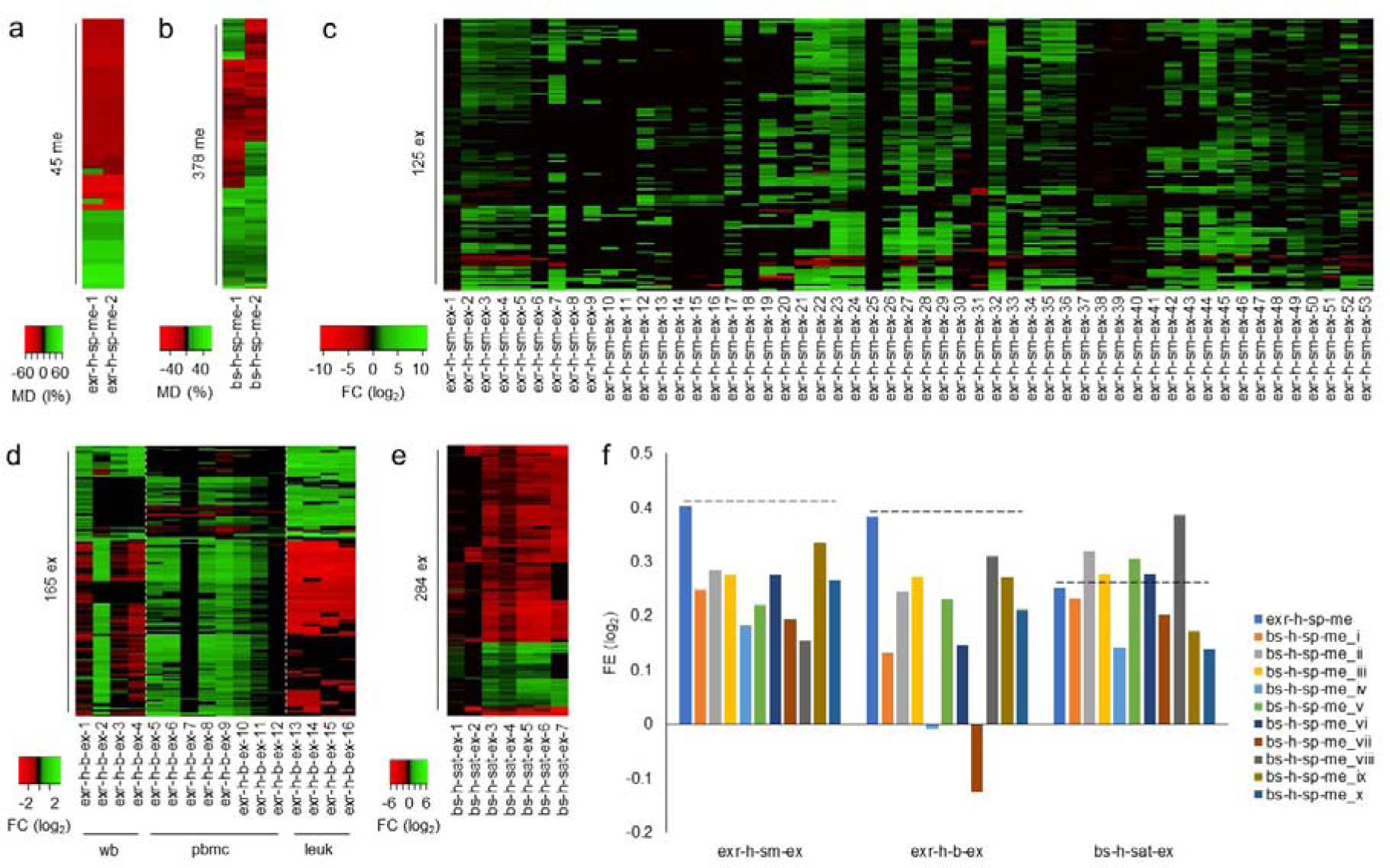
Gene set overlap between human sperm DMGs and human somatic DEmRNAs. Heatmaps showing methylation difference in exercise (**a**) and bariatric surgery (**b**) associated sperm DMGs at two time points, and showing fold change of most common DEmRNAs in skeletal muscle (**c**) and blood (**d**) following exercise, and in (**e**) subcutaneous adipose tissue following bariatric surgery. The number of DMGs and DEmRNAs in the heatmaps are indicated. b, blood; bs, bariatric surgery; ex, differentially expression; exr, exercise; FC, fold change; h, human; leuk, leukocytes; MD, methylation difference; me, differential methylation; pbmc, peripheral blood mononuclear cells; sat, subcutaneous adipose tissue; sm, skeletal muscle; sp, sperm; wb, whole blood. Individual gene sets are unique and intuitively labeled, with intervention, species, tissue, data type, and sample category multiplicity indicated in that order and separated by a dash. (**f**) Bar graph showing fold enrichment of pooled sets of skeletal muscle (exr-h-sm-ex), blood (exr-h-b-ex), and subcutaneous adipose tissue (bs-h-sat-ex) DEmRNAs in the pooled set of exercise associated sperm DMGs (exr-h-sp-me) and size-matched non-overlapping subsets of pooled bariatric surgery associated sperm DMGs (bs-h-sp-me_i to x). FE, fold enrichment. Full sets of DMGs and DEmRNAs, along with gene set identifiers used here and other details, are provided in **Supplementary Table 1**. The source data (a-f) are provided as **Source Data file 1**.

### Overlap between human exercise sperm DMGs and animal exercise somatic DEmRNAs

The exercise and bariatric surgery associated human sperm DMGs were next tested for overlap with exercise associated DEmRNAs in animal models. The animal data included DEmRNAs in skeletal muscle in rats and mice, and in larval body wall muscles and whole fly homogenate in *Drosophila*. As species-specific clustering of DEmRNAs showed an overall trend for directional similarities in gene expression regulation across samples (**Fig. 2a-c**), the DEmRNAs belonging to each of these species were, as in human alone analysis above, separately pooled for overlap test. Notably, the human exercise sperm DMGs, in comparison to human bariatric surgery sperm DMGs, showed unambiguously higher enrichment of animal exercise somatic DEmRNAs in rat, mouse, and fly models alike (**Fig. 2d**). Cumulatively, these results demonstrated that exercise human sperm DMGs represent exercise response pathways conserved across species.

**Fig. 2.**
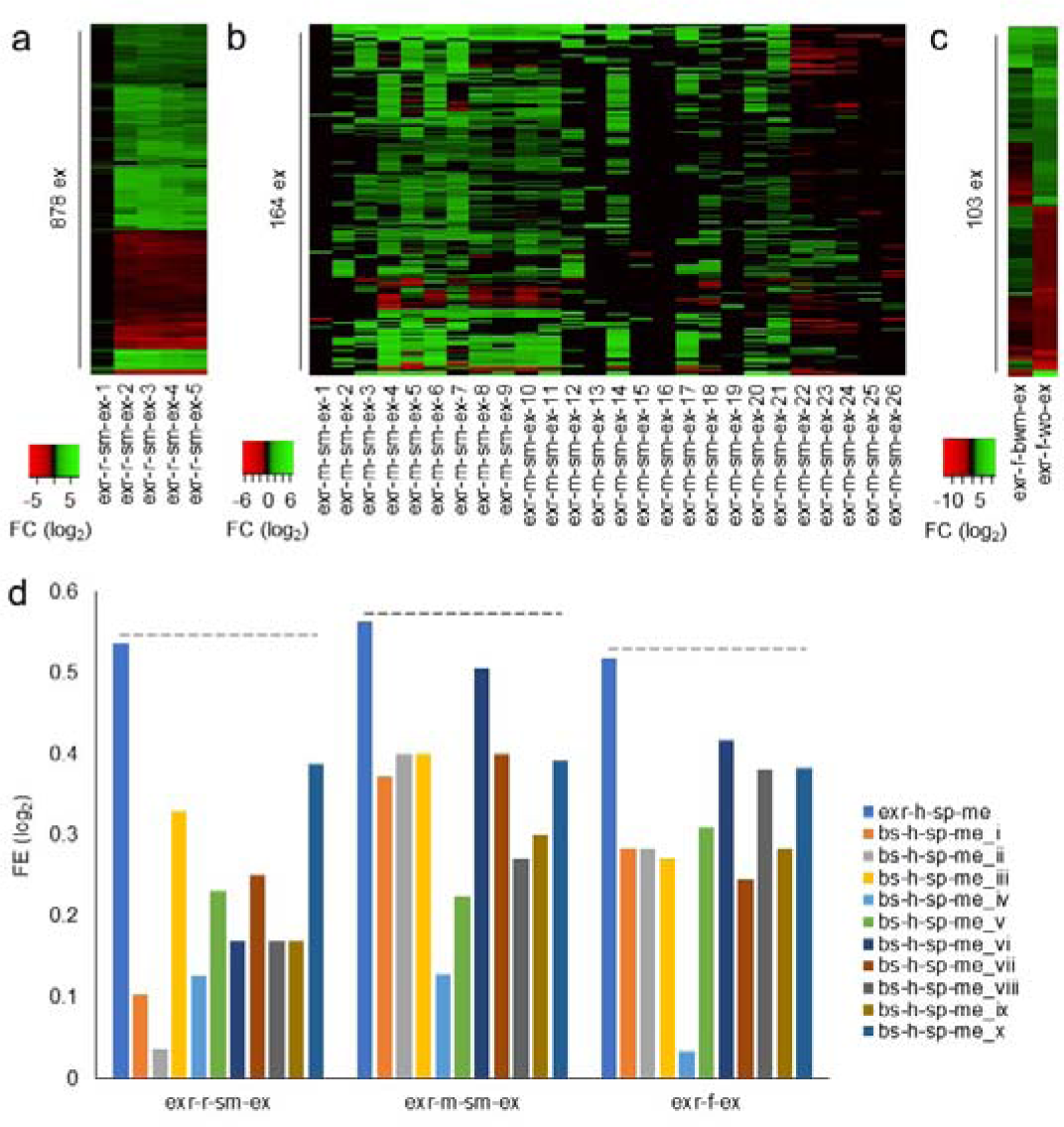
Gene set overlap between human sperm DMGs and animal somatic DEmRNAs. Heatmaps showing fold change of most common exercise associated DEmRNAs in rat (**a**) and mouse (**b**) skeletal muscle, and of (**c**) exercise associated DEmRNAs in both *Drosophila* larval body wall muscles and whole fly samples. r, rat; m, mouse; f, fly; bwm, larval body wall muscles; wo, whole organism. Labeling scheme of individual samples, and abbreviations as explained in Fig. 1. (**d**) Bar graph showing fold enrichment of pooled sets of rat (exr-r-sm-ex) and mouse (exr-m-sm-ex) skeletal muscle, and of *Drosophila* larval body wall muscles and whole fly (exr-f-ex) DEmRNAs in the pooled sets of exercise associated sperm DMGs and size-matched non-overlapping subsets of pooled bariatric surgery associated sperm DMGs. Other details as mentioned in Fig. 1. Full sets of DMGs and DEmRNAs, along with gene set identifiers used here and other details, are provided in **Supplementary Table 1**. The source data (a-d) are provided as **Source Data file 2**.

### Gene ontology enrichment in exercise sperm DMGs and exercise somatic DEmRNAs

Next, all exercise associated human and animal gene set pools used above for overlap analysis were subjected to gene ontology analysis for enrichment of biological process terms. Notably, among various commonly enriched terms, muscle structure development showed highest enrichment in sperm DMGs, with the term also showing high enrichment in human, rat, and mouse skeletal muscle DEmRNAs, and *Drosophila* larval body wall muscles and whole fly DEmRNAs combined (**Fig. 3a**). Intuitively, human blood DEmRNAs alone did not show enrichment of muscle structure development. A survey of other “muscle” containing terms enriched in gene sets revealed several of these terms in both sperm DMGs and somatic, except blood, DEmRNAs (**Fig. 3b**). These results further supported the evidence that exercise associated sperm DMGs relate to exercise biology.

**Fig. 3.**
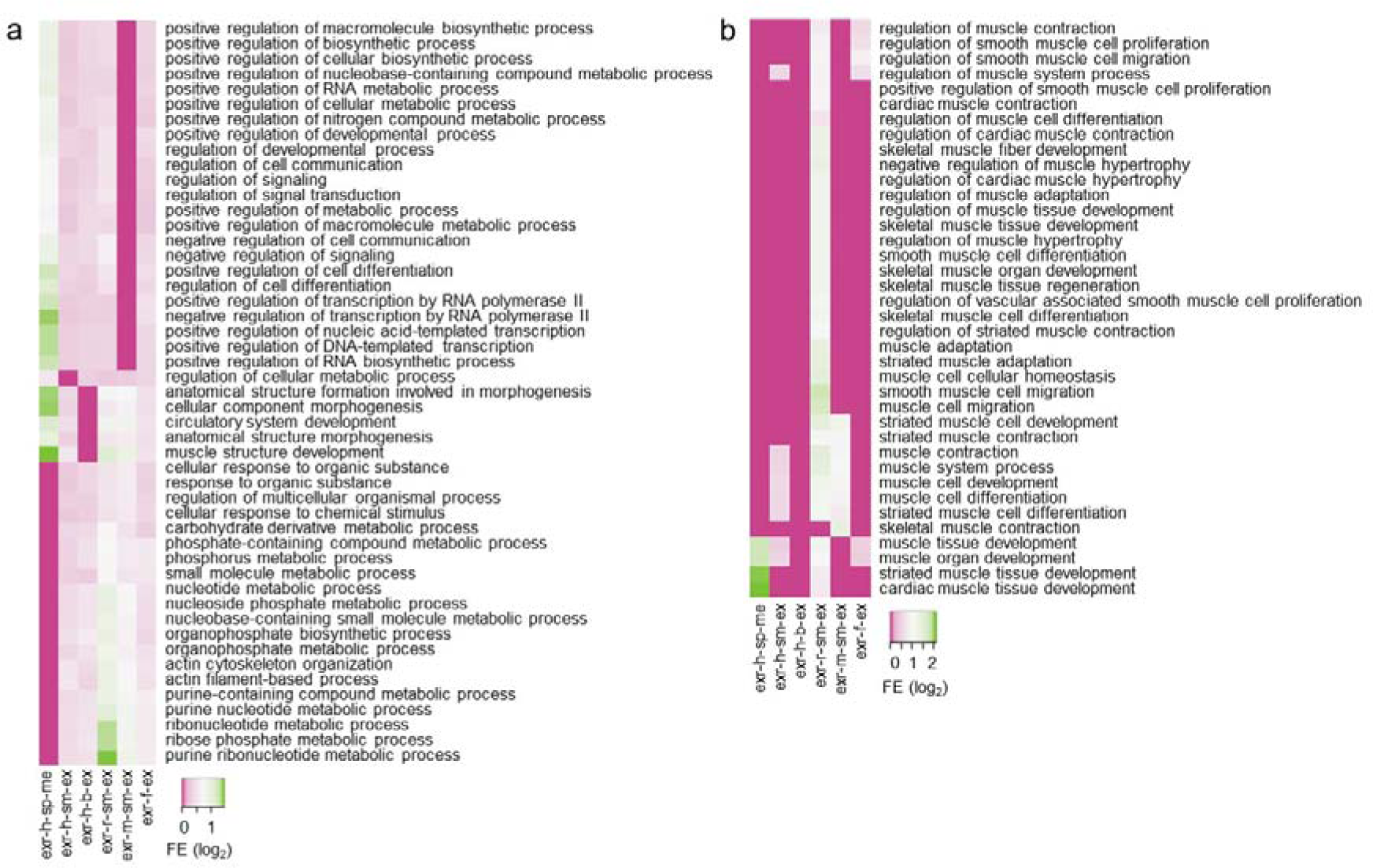
Gene ontology enrichment in human and animal model DMGs and/or DEmRNAs. (**a**) Heatmap showing fold enrichment of biological process terms most commonly overrepresented among pooled sets of exercise associated human sperm DMGs (exr-h-sp-me), human skeletal muscle (exr-h-sm-ex) and blood (exr-h-b-ex) DEmRNAs, rat (exr-r-sm-ex) and mouse (exr-m-sm-ex) skeletal muscle DEmRNAs, and *Drosophila* (exr-f-ex) DEmRNAs. (**b**) Heatmap showing fold enrichment of rest of the overrepresented “muscle” containing terms. The full list of enriched terms is provided in **Supplementary Table 2**. Other details as mentioned in Fig. 1 and 2.

### Directionality of human exercise sperm DMGs and human exercise somatic DEmRNAs

The exercise sperm DMGs and exercise somatic DEmRNAs in humans were next compared in terms of regulatory directionality. For this, the DMGs common to both individual sperm samples (**Fig. 1a**) were separated in two groups, promoter and non-promoter, with the latter group representing intron, exon, downstream, distal intergenic, and 5’ UTR combined. The promoter and non-promoter DMGs were then aligned with all the individual exercise and bariatric surgery somatic DEmRNA samples (**Fig. 1c-e**), for comparing directionality of regulation in DMGs with that in DEmRNAs (**Fig. 4a**). Startlingly, chi-square goodness of fit test showed that the observed counts of DEmRNA occurrences showing opposite of DMG regulatory directionality were, assuming equal chance for same and opposite regulation, significantly greater than the expected counts in the exercise, not bariatric surgery, group, for promoter, not non-promoter, set (**Fig. 4b**). These findings crucially supported the hypothesis that exercise induced somatic gene expression changes are encoded in sperm epigenome.

**Fig. 4.**
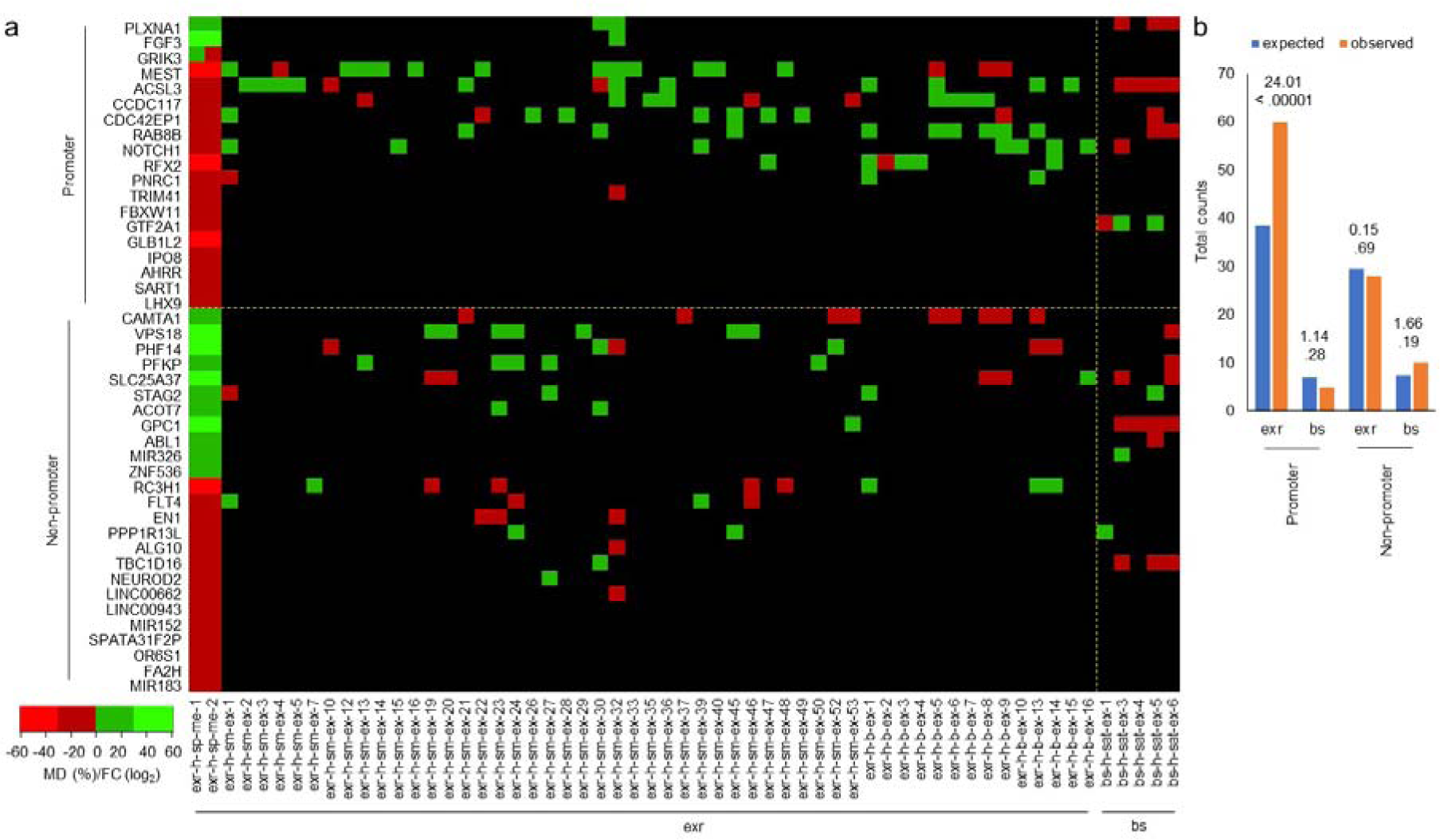
Directionality of human sperm promoter DMGs and somatic tissue DEmRNAs. (**a**) Heatmap of exercise associated human sperm promoter and non-promoter DMGs (methylation difference) that are common for the two time points, shown along with corresponding DEmRNAs (fold change) in human exercise associated skeletal muscle and blood, and bariatric surgery associated subcutaneous adipose tissue. DEmRNA samples present in Fig. 1c-e but missing here indicate no DMG common DEmRNA therein. A reduced number of heatmap shades was used to discretize regulation. (**b**) Bar graph showing expected and observed counts of DEmRNA occurrences with regulatory direction inverse of DMGs, shown in the heatmap (a). Chi-square goodness of fit and corresponding *p* values have been shown above the bars. Other details as mentioned in Fig. 1. Full sets of DMGs and DEmRNAs, along with gene set identifiers used here and other details, are provided in **Supplementary Table 1**. The source data (a, b) are provided as **Source Data file 3**.

### Exercise sperm promoter DMGs, miRNA targets, and somatic DEmiRNAs in humans

In the last step of the analysis, miRNA target enrichment in human exercise sperm promoter DMGs was examined first. In this, highest enrichment was found for hsa-miR-181 family miRNAs (**Fig. 5a**). It was next examined if this family of miRNAs are differentially expressed in soma following exercise. Interestingly, these miRNAs were frequently represented among exercise associated DEmiRNAs in blood cells and plasma (**Fig. 5b**). However, while these exercise somatic DEmiRNAs showed upregulation (**Fig. 5b**), their targets among the exercise sperm promoter DMGs (**Fig. 4a**), namely, *CCDC117*, *GRIK3*, *IPO8*, *LHX9*, *PNRC1*, *RAB8B*, and *RFX2*, were mostly hypomethylated, with the corresponding exercise somatic DEmRNAs (**Fig. 4a**), when present, mostly upregulated. Given that miRNAs usually repress gene expression and hence a negative correlation is expected between miRNA and mRNA, the positive correlation found above seems counterintuitive. However, several instances of positive correlations are also known, with the bidirectionality explained in terms of complex gene regulatory networks (Diaz *et al*., 2015; Laxman *et al*., 2015). Finally, gene ontology analysis of hsa-miR-181 family targets showed enrichment of various biological process terms, with several muscle related ones showing high enrichment (**Fig. 5c**). The hsa-miR-181 family miRNAs are associated with a diversity of exercise responses. For example, plasma hsa-miR-181c levels is a predictive marker of exercise training response in heart failure patients (Gevaert *et al*., 2021). Also, circulating levels of hsa-miR-181c in peripheral blood has been associated with exercise capacity and ventilatory inefficiency in acute myocardial infarction (Miyazawa *et al*., 2021). Similarly, expression levels of miRNAs including hsa-miR-181a in extracellular vesicles in plasma has been correlated with aerobic fitness in young men (Guescini *et al*., 2015). Likewise, exercise training has been found to cause increased miR-181a expression levels in testicular tissue in opioid treated rats (Hadi *et al*., 2022). Cumulatively, the above results consolidated the evidence that exercise sperm DMGs are functionally related to exercise biology.

**Fig. 5.**
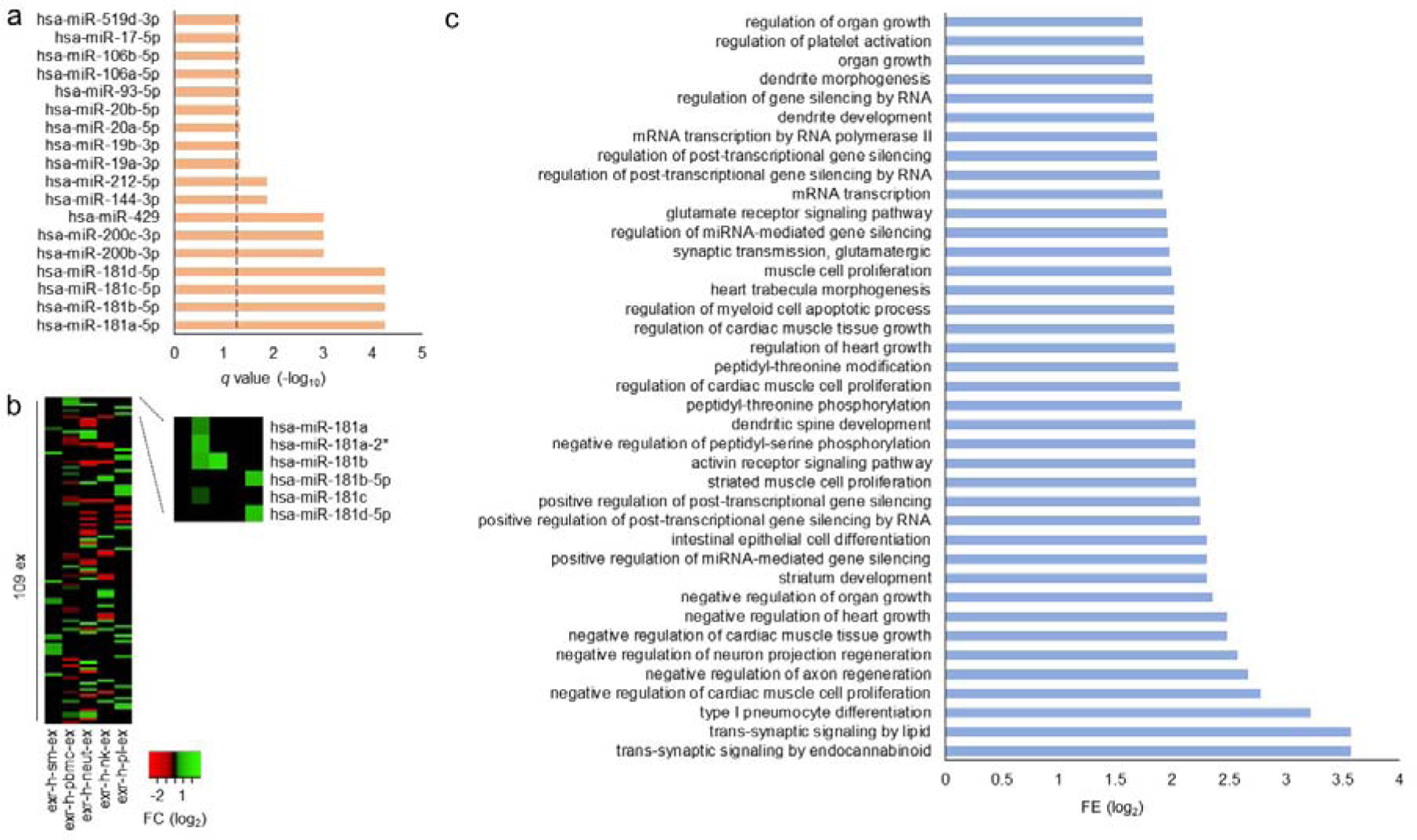
Enrichment of miRNA targets in human sperm promoter DMGs. (**a**) Bar graph showing Bonferroni *q* value of fold enrichment for targets of various miRNAs in DMGs. The dashed line indicates cutoff for statistical significance. (**b**) Heatmap showing fold change of DEmiRNAs most commonly identified among various tissues in human volunteers following exercise. The hsa-miR-181 family miRNAs are blown-up in the inset. neut, neutrophils; nk, natural killer cells; pl, plasma. (**c**) Bar graph showing fold enrichment of most overrepresented biological process terms in hsa-miR-181 family targets. The complete list of enrichment terms is given in **Supplementary Table 3**. Other details as mentioned in Fig. 1. Full sets of DEmiRNAs, along with miRNA set identifiers used here and other details, are provided in **Supplementary Table 1**. The source data (b) are provided as **Source Data file 4**.

Exercise associated alterations in the levels of circulating hsa-miR-181 family miRNAs in the genome level datasets analysed, and reported in the other studies including that on extracellular vesicle (Gevaert *et al*., 2021; Guescini *et al*., 2015; Miyazawa *et al*., 2021), together with the evidence that exercise influences the levels of one of the family members in testicular tissue in rats (Hadi *et al*., 2022), call for a tempting speculation. Given previous bioinformatic evidence implicating extracellular circulating miRNAs as potential carriers of soma to germline information transfer in epigenetic inheritance (Sharma, 2014), and its subsequent experimental support (Gapp *et al*., 2014; Sharma *et al*., 2016; Chen *et al*., 2016), the above associations of hsa-miR-181 family miRNAs in exercise seems tantalizing. It is tempting to speculate that these miRNAs may mediate soma to germline communication in inheritance of exercise induced traits.

The analysis presented here shows that transcriptomic response to physical exercise in soma is functionally represented in the germline epigenome in humans, thus elevating the plausibility of inheritance of induced traits in our species. Future cross-generational human studies may provide more definitive evidence. With exercise being a common healthy practice recommended by leading organizations, its effects in the offspring beneficial in rodents (Kusuyama *et al*., 2020; Vieira de Sousa Neto *et al*., 2021), sperm and peripheral blood samples convenient, and, now, inheritance of exercise induced traits in humans supported, an ethically ideal opportunity has arisen to undertake such studies. In the latter, sperm from prospective fathers in exercise and non-exercise cohorts can be first used to confirm expected exercise induced sperm DNA methylation changes, and upon confirmation, the studies can be extended and offspring peripheral blood tested for expected exercise induced gene expression changes. Such hypothesis testing human studies are all the more warranted considering that epigenetic inheritance could offer an explanation for missing causality and heritability of complex diseases (Slatkin, 2009; Furrow *et al*., 2011; Danchin *et al*., 2011; Trerotola *et al*., 2015) and epigenetic drugs could provide a means for reversing disease associated epigenetic states (Heerboth *et al*., 2014; Kronfol *et al*., 2017; Rittiner *et al*., 2022), as also in view of the ethical concerns of epigenetic inheritance (Juengst *et al*., 2014; Sasaki-Honda *et al*., 2023).

## Supporting information

Supplementary Table 1. Data compendium

Supplementary Table 2. Gene ontology enrichment in exercise associated gene sets

Supplementary Table 3. Gene ontology enrichment in has-miR-181 family targets

Source Data file 1. Source data for Fig. 1a-e

Source Data file 2. Source data for Fig. 2a-d

Source Data file 3. Source data for Fig. 4a, b

Source Data file 4. Source data for Fig. 5b

## Declaration of Interest Statement

The author declares no conflict of interest.

## Supplemental Materials

**Supplementary Table 1.** Data compendium

**Supplementary Table 2.** Gene ontology enrichment in exercise associated gene sets

**Supplementary Table 3.** Gene ontology enrichment in has-miR-181 family targets

**Source Data file 1.** Source data for Fig. 1a-e

**Source Data file 2.** Source data for Fig. 2a-d

**Source Data file 3.** Source data for Fig. 4a, b

**Source Data file 4.** Source data for Fig. 5b

## Notes

### Competing Interest Statement

The authors have declared no competing interest.

### Summary of Updates

The manuscript has been revised for brevity in text and results, enhanced readability and accessibility of the content, facilitated reproducibility of the results with newly added supplementary files containing the source data for figures, and removing typographical and formatting errors in the supplementary tables, with data and figures remaining completely intact.

